# ChromMovie: A Molecular Dynamics Approach for Simultaneous Modeling of Chromatin Conformation Changes from Multiple Single-Cell Hi-C Maps

**DOI:** 10.1101/2025.05.16.654550

**Authors:** Krzysztof H. Banecki, Haoxi Chai, Yijun Ruan, Dariusz Plewczynski

## Abstract

The development of 3C-based techniques for analyzing three-dimensional chromatin structure dynamics has driven significant interest in computational methods for 3D chromatin reconstruction. In particular, models based on Hi-C and its single-cell variants, such as scHi-C, have gained widespread popularity. Current approaches for reconstructing the chromatin structure from scHi-C data typically operate by processing one scHi-C map at a time, generating a corresponding 3D chromatin structure as output. Here, we introduce an alternative approach to the whole genome 3D chromatin structure reconstruction that builds upon existing methods while incorporating the broader context of dynamic cellular processes, such as the cell cycle or cell maturation. Our approach integrates scHi-C contact data with single-cell trajectory information and is based on applying simultaneous modeling of a number of cells ordered along the progression of a given cellular process. The approach is able to successfully recreate known nuclear structures while simultaneously achieving smooth, continuous changes in chromatin structure throughout the cell cycle trajectory. Although both Hi-C-based chromatin reconstruction and cellular trajectory inference are well-developed fields, little effort has been made to bridge the gap between them. To address this, we present ChromMovie, a comprehensive molecular dynamics framework for modeling 3D chromatin structure changes in the context of cellular trajectories. To our knowledge, no existing method effectively leverages both the variability of single-cell Hi-C data and explicit information from estimated cellular trajectories, such as cell cycle progression, to improve chromatin structure reconstruction.

## Introduction

The modeling of three-dimensional (3D) chromatin structure has garnered significant attention in recent years, largely due to the development of advanced techniques that provide deeper insight into chromatin organization. Methods for analyzing 3D genome organization can be broadly categorized into microscopic imaging-based approaches, sequencing-based techniques, and their accompanying computational models [1]. While microscopic methods, such as super-resolution imaging combined with fluorescence in situ hybridization (FISH) [2], 3D-EMISH [3] or iPALM [4], have been applied to 3D chromatin structure reconstruction [5], sequencing-based computational approaches have been far more widely adopted. These methods have led to a diverse range of computational strategies for chromatin modeling [6, 7] and have been extensively reviewed in the literature [8,9,10,11,12].

## Single-cell Hi-C

Several sequencing-based methods have been developed to provide genome-wide information on chromatin conformation. These include ligation-based techniques such as Hi-C [13], HiChIP [14], Micro-C [15] or ChIA-PET [16], as well as non-ligation approaches such as SPRITE [17], ChIA-Drop [18] or GAM [19]. Among these, Hi-C and its derivatives have been the most widely used for chromatin structure reconstruction, offering invaluable insights into chromatin conformations across genomic scales [13, 20, 21]. However, reconstructing 3D chromatin structures from Hi-C data presents challenges, for example in resolving ambiguous knotting patterns [22] or in addressing the ergodicity of interaction frequencies [23]. To overcome at least some of these limitations, single-cell and single-nucleus Hi-C techniques have been developed [24, 25, 26, 27], enabling novel insights into chromatin conformation heterogeneity at the single-cell level. For a detailed overview of published single-cell and single-nucleus datasets, see [28, 29, 30, 31].

While Hi-C data is commonly used for chromatin structure reconstruction, single-cell approaches offer distinct advantages by bridging the gap between bulk Hi-C’s averaged view and microscopy-based studies of single-cell conformations [24]. Several methods have been developed for reconstructing the 3D chromatin structure from single-cell data [24, 32, 33, 34, 27, 35, 36, 37, 38, 39, 40, 41] which were reviewed in [42], and new ones are constantly being developed [43]. Despite their methodological diversity, these approaches share key similarities. Notably, most models reconstruct chromatin structure using a single scHi-C map at a time, potentially overlooking contextual information from other cells. Incorporating this broader context could improve reconstruction accuracy—for example, by integrating bulk Hi-C data, as demonstrated in [35]. Another method of using this context information indirectly is by applying imputation techniques for which several notable examples include Higashi [44], Fast-Higashi [45], ScHiCEDRN [46], HiC-SGL [47], ScHiCAtt [48] or HiCENT [49]. These methods, and other general deep learning models developed for Hi-C resolution enhancement [50, 51], can be used in tandem with chromatin reconstruction algorithms to enrich the resulting structures. However, most do not explicitly model structural changes along a continuous cellular trajectory or process of interest.

### Trajectory inference methods

Broadening our perspective beyond Hi-C studies within cellular genomics, we find significant advances in single-cell analysis. For instance, single-cell RNA sequencing (scRNA-seq) has revolutionized transcriptomics by offering advantages over traditional bulk analysis [52,53]. Single-cell transcriptomics has provided key insights into the inherent heterogeneity of dynamic cellular processes, including cell differentiation, maturation, activation, response to stimuli, and cell cycle progression. To study the transitions cells undergo during these dynamic processes, numerous computational methodologies—collectively known as trajectory inference (TI) methods—have been developed and extensively reviewed [54,55,56], alongside methods focused more specifically on single-cell clustering [57]. The field continues to expand, with dozens of TI methods already established and new ones continually emerging [58,59,60,61,62,63,64].

While RNA-seq-based TI methods have been widely applied to studying cell cycle trajectories, cell maturation, and differentiation, much less effort has been made to integrate these approaches with single-cell Hi-C data and, consequently, 3D chromatin modeling. One of the first attempts to reconstruct a cell cycle trajectory using scHi-C data was by Nagano et al. (2017) [65], who introduced the ‘repli-score’ to estimate pseudo-trajectories of cell cycle progression. Subsequently, methods such as CIRCLET [66] or scHiCPTR were developed with similar objectives. Additionally, several approaches have been proposed for clustering or classification of scHi-C data, particularly for identifying cell cycle phases. Notable recent examples include scHiCluster [68], Kim et al. (2020) [69], scHiCyclePred [70] or scHiClassifier [71].

### Bridging the gap

Beyond trajectory inference and clustering, other key areas of single-cell Hi-C analysis include embedding, imputation, and denoising, all of which remain active research topics. Most scHi-C analyses begin with dimensionality reduction and embedding (for a review of embedding methods in scHi-C, see [72]). These embeddings serve as a foundation for subsequent analyses, such as imputation [44, 45] and denoising [73, 74], and are also valuable starting points for reconstructing cell cycle trajectories and other dynamic cellular processes.

We propose that integrating cellular trajectory information obtained from TI methods adapted for scHi-C could significantly enhance 3D chromatin reconstruction. To date, no existing method explicitly incorporates trajectory data into chromatin structure reconstruction from scHi-C. Most reconstruction methods are based on Hi-C, with only a few attempting to integrate microscopy data [75] or additional genomic datasets such as CHIP-seq or RNA-seq [76, 77]. While an increasing number of approaches incorporate single-cell or single-nucleus Hi-C data [42], they generally treat each scHi-C map independently, thereby losing valuable information embedded in cellular trajectories.

Here, we introduce *ChromMovie*, the first 3D chromatin conformation modeling method — to our knowledge — that explicitly incorporates cellular trajectory information, such as cell cycle or maturation dynamics. ChromMovie uses an OpenMM framework [78] and builds upon widely used molecular approaches in scHi-C-based chromatin structure reconstruction [42] by modeling multiple scHi-C maps simultaneously and leveraging their ordering, as determined by a TI method or the scHi-C experiment itself (see Fig. 1A). The method incorporates the knowledge about cell order by adding meta-restraints to the molecular dynamics simulation which itself is based on a previous methods for 3D chromatin structure reconstruction from scHi-C data such as NucDynamics [27]. In this way the simulation of a 3D chromatin structure from a particular scHi-C map is influenced by neighboring scHi-C maps within the dataset, with neighbors defined as the immediate predecessor and successor in the trajectory order. Rather than treating each scHi-C map independently, this approach allows chromatin contacts from adjacent pseudo-time points to influence structure formation, capturing continuity along the trajectory. By incorporating this temporal context, ChromMovie enhances robustness against the inherent noise in scHi-C data and serves as an implicit normalization strategy. This prevents reconstructed chromatin structures at any pseudo-time point from becoming trapped in local optima of molecular dynamics simulations, leading to more biologically meaningful results.

**Fig. 1.**
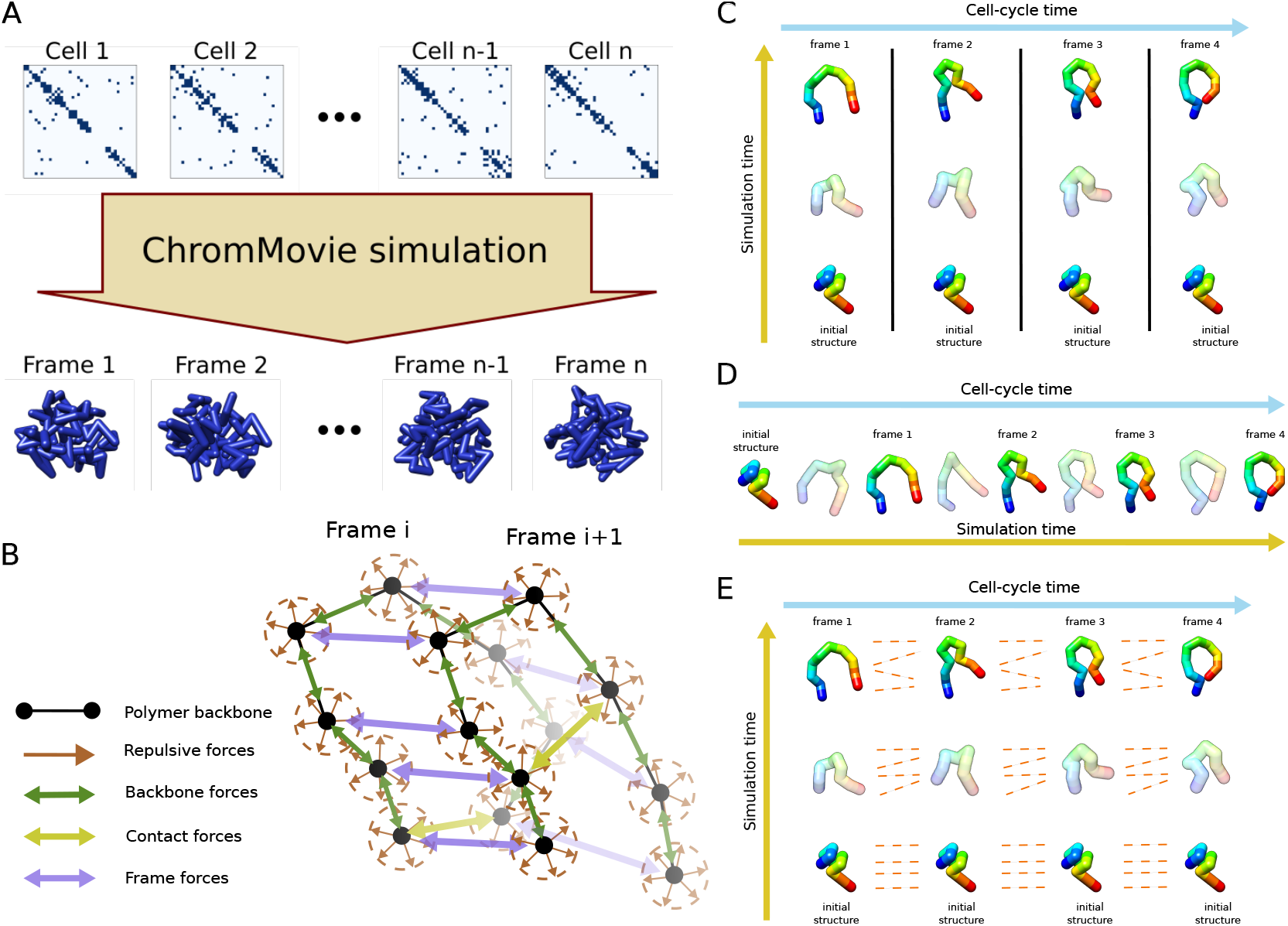
(A) Overview of the ChromMovie algorithm. The algorithm takes an ordered list of scHi-C maps representing different time points of a cellular process. It then simulates all time points simultaneously, generating an ordered list of 3D structures as the result. (B) The four main types of forces used in the ChromMovie simulation: repulsive forces, backbone forces, contact forces, and frame forces. (C, D, E) Three approaches to modeling multiple ordered scHi-C maps: independent modeling (C), sequential modeling (D), and parallel simultaneous modeling (E). ChromMovie implements a form of simultaneous modeling, which, to our knowledge, is the first of its kind in the field of chromatin modeling.

## Method description

We identified three fundamental approaches for molecular dynamics (MD) simulations of chromatin structures based on multiple scHi-C maps, which are graphically illustrated in Fig. 1C, D and E.

1. Independent Approach – Each scHi-C map is processed separately using any existing chromatin reconstruction method, without incorporating information from other maps.
2. Sequential Approach – Modeling begins with the first scHi-C map in the presumed order. The output of each simulation serves as the starting structure for the next, incorporating constraints from the subsequent scHi-C map.
3. Semi-Dependent Approach – Each scHi-C map is modeled with its own set of constraints, but additional restraints ensure that structures from consecutive maps remain spatially aligned by applying forces that link corresponding chromatin beads.

The independent approach is highly susceptible to overfitting due to noise in individual scHi-C maps. This limitation makes the sequential and semi-dependent approaches potentially more advantageous, as they incorporate information from neighboring matrices, providing a more coherent and biologically meaningful reconstruction of chromatin dynamics. The two alternative approaches aim to reduce the risk of overfitting while incorporating contextual information from neighboring scHi-C maps. The sequential approach achieves this by explicitly integrating structural information from previous frames, using each reconstructed chromatin structure as the starting point for the next phase of modeling. As a result, although indirectly, information from all preceding frames influences the reconstruction of a given frame.

However, while the independent approach disregards both preceding and subsequent frames when modeling single-cell chromatin structures, the sequential approach also fails to incorporate information from later frames that could further refine the reconstruction. To fully leverage the available information encoded in cellular trajectories, we developed a third strategy: the simultaneous approach (Fig. 1E). By introducing connections between consecutive frames along the cellular trajectory, this approach utilizes information from both earlier and later pseudo-time points. ChromMovie is the first method designed to integrate this temporal context, enhancing the accuracy of single-cell chromatin conformation reconstruction while capturing its dynamic changes throughout a cellular process.

### Force field

As outlined in a recent review [42], most existing methods for 3D chromatin structure reconstruction define their potential function using a few fundamental components. These typically include repulsive forces applied to all possible pairs of chromatin beads, as well as attractive forces. The attractive forces are primarily applied to (i) the chromatin polymer backbone, ensuring connectivity between consecutive beads along the chromatin chain, and (ii) interactions between beads linked by scHi-C contacts. Our simulation follows this general force field framework while introducing an additional fourth type of force (Fig. 1B). This force is designed to connect corresponding chromatin beads across different pseudo-time frames, capturing the continuity of chromatin dynamics during processes such as the cell cycle or cell maturation.

Let *n* be the number of pseudo-time frames to be modeled and *m* the number of chromatin beads in each frame. The input to our simulation consists of an ordered list of *n* binary matrices H*i* each of size *m* × *m*, encoding the loci connected by single-cell contacts. Let **v**_*i*,*j*_ = (*x*_*i*,*j*_ , *y*_*i*,*j*_ , *z*_*i*,*j*_) denote the three-dimensional position of the *j*-th chromatin bead of the *i*-th pseudo-time frame, corresponding to the single-cell contact matrix H*i*. Furthermore, let *d*(**v**_1_, **v**_2_) represent the Euclidean distance between the vectors **v**_1_ and **v**_2_.

With these definitions, the four main components of our general potential function, in its simplest form, are as follows:

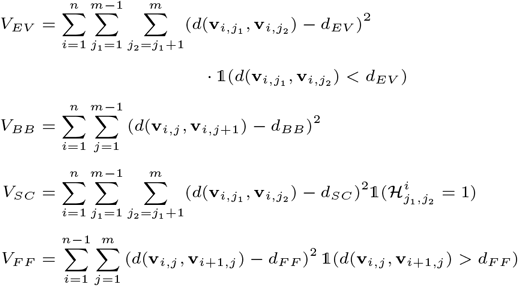

with the general potential *V* taking the form:

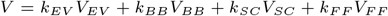

where *k*_*EV*_ , *k*_*BB*_, *k*_*SC*_ and *k*_*F F*_ are all user-specified coefficients controlling the relative importance of each of the four force field components. Similarly, the distances *d*_*EV*_ , *d*_*BB*_, *d*_*SC*_ and *d*_*F F*_ are user-specified “optimal” distances for each of these forces.

The equations above represent the most basic form of the ChromMovie force field, featuring harmonic or harmonic-like potentials applied to the four main components of the force field. However, drawing from a wide range of established methods for 3D chromatin reconstruction from scHi-C data, we incorporated several enhancements into these basic force components.

First, the strict harmonic potential is often considered too stringent, as it can strongly influence simulation dynamics [36]. This potential function rapidly increases to large values with even small deviations from the optimal target distance *d*_*opt*_, which could lead to significant conflicts with other forces in the system and make it harder for the simulation to reach a satisfactory potential optimum. To address this issue, ChromMovie incorporates several effective strategies from previous studies. One such strategy is replacing parts of the quadratic function with linear sections. This is done by introducing a “flat bottom” potential between 0.8 · *d*_*opt*_ and 1.2 · *d*_*opt*_ (similarly to [27]), or approximating the quadratic function with a linear function for larger distances (e.g. [24] or [27]). These modifications create a less stringent restraint, allowing for a wider range of possible distances and a more gradual increase in energy when the distance requirement is not met during the simulation. Finally, if an upper limit on the force is required, a Gaussian potential can also be used (as in [36]). In ChromMovie, the choice between using Gaussian, harmonic, or harmonic with linear end potentials is left to the user.

Following the example of several previous methods (e.g. [79, 27, 35, 38, 41]), ChromMovie is also a hierarchical model. Starting at the lowest resolution, the simulation is performed at each resolution specified by the user. After each round of molecular dynamics (MD) simulation, the resulting structure is interpolated so that the number of beads in the structure matches the corresponding resolution. Naturally, the optimal distances for all forces must be rescaled after each resolution change and interpolation. In this case, we assumed that the optimal distances provided by the user represent the distances between beads at the final (and lowest) resolution. The optimal distances for intermediate resolutions are determined using a power-law relationship with a coefficient of 1*/*3 to ensure a smooth transition between resolutions.

The final improvement pertains to the handling of single-cell contact information. An enhancement to the classical MD chromatin simulation involves tuning the optimal distance for single-cell contacts according to the number of contacts in the corresponding scHi-C map entry. This allows us to effectively utilize the number of contacts between each pair of beads without resorting to the binarization of contact matrices, as seen in some previous methods [34]. Following the approach outlined in [27], we set the optimal distance for contact loci to be proportional to *c*^−1*/*3^ where *c* is the number of contacts between two chromatin beads. Additionally, ChromMovie offers an optional feature that discards “violated” contacts—those for which the 3D distance fails to converge to the desired range within the simulation time.

### Simulation

The MD simulation in ChromMovie is performed using a standard Langevin integrator. The initial structure for the simulation is a self-avoiding random walk. To avoid issues with initial discordance and violations of the forces that connect consecutive cellular beads, the same initial structure is applied to all frames. The simulation is performed hierarchically, meaning that after the simulation is over for a particular resolution, the structure beads are interpolated in preparation for the next phase of the simulation with a finer resolution. Each simulation phase is parallelized to enhance computational time and take advantage of GPU computing.

Avoiding local minima in simulations is a common challenge, particularly in MD simulations and optimization tasks. In our algorithm, the goal of a successful cellular trajectory reconstruction is to satisfy the restraints on chromatin backbone closeness, single-cell contacts, and the forces keeping consecutive cellular frames together. The force counteracting these restraints is the excluded volume (EV) force, which is applied to all structure beads within a given frame (but not between frames).

To enhance the algorithm’s ability to reach the potential minimum without violating the imposed restraints, some models (e.g. [41]) gradually reinforce the force coefficients throughout the simulation. Based on this idea, we have found that adjusting coefficients during simulation may help the model satisfy single-cell restraints (see Results section). ChromMovie allows users to enable or disable gradual reinforcement of the coefficients across all four force field components. The coefficient reinforcement scheme is similar to the one used in [27], and is based on arctan function. During the simulation, the initial coefficient *k*_*XX*,*init*_ specified by the user is multiplied by this function, starting with a small coefficient and gradually reaching the full value of *k*_*XX*,*init*_ towards the end, as described below:

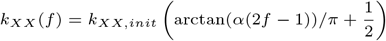

Here the *f* is the elapsed proportion of simulation and *α* is the parameter controlling the steepness of the transition. In our model we used *α* = 10 same as in [27].

## Results

To evaluate the performance of our model, we conducted several analyses using both in silico examples and real-life single-cell contact data, incorporating cellular trajectory context derived from [80]. The in silico studies enabled us to fine-tune the hyperparameters of ChromMovie, while the real-life data helped us demonstrate the algorithm’s effectiveness in studying changes in single-cell chromatin conformation.

### Benchmarking on in silico models

To validate our model, we first tested it on an artificial in silico model. We chose a process involving the contraction of a ring-like zigzag structure (see Fig. 2A) with a constant distance between consecutive beads. Based on this ‘ground truth’ structure, we generated a number of contacts for each frame, proportional to the 3D distance between loci, which were then used as input for ChromMovie. This allowed us to assess how well the model reconstructed the original structure by comparing it to the modeled structures using the Root Mean Square Deviation (RMSD) measure.

**Fig. 2.**
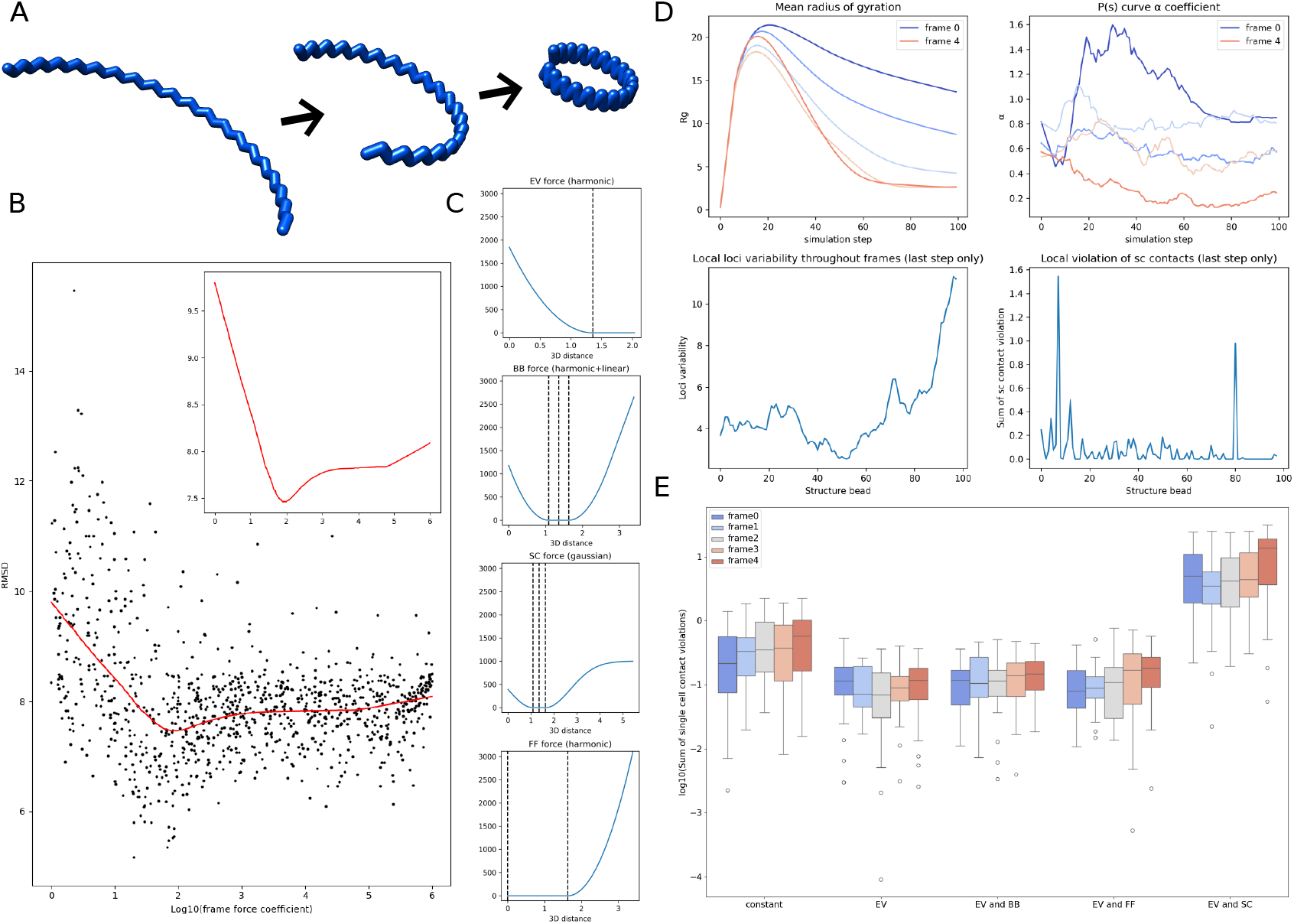
ChromMovie in silico analysis. (A) In silico example of a ring-forming contracting zigzag structure used for initial validation. (B) Results of 1, 000 runs of the ChromMovie simulation on the in silico model. Root Mean Square Deviation (RMSD) was used to compare with the original structure. The top-right corner shows a zoomed-in view of the fitted Loess curve, indicating a clear minimum. (C) Examples of the force potentials used in ChromMovie. From top to bottom: harmonic excluded volume (EV) potential, harmonic backbone (BB) potential with linear approximation for larger distances, Gaussian single-cell (SC) potential, and harmonic frame-force (FF) potential. (D) Selected diagnostic plots from the ChromMovie reporter. The mean radius of gyration indicates greater compaction for frames closer to the end, consistent with the original structure. *P* (*s*) curve allows for analysis of the contact frequency distribution. Local loci variability helps visualize regions of the structure that experience the greatest changes during the studied cellular process. (E) Single-cell contact violations across different ChromMovie runs, comparing cases with the gradual increase of the respective force coefficient turned on or kept constant.

We initially used this approach to validate the core concept of ChromMovie, which is to enhance general MD chromatin reconstruction by using connections between frames. For this, we performed 1, 000 runs of ChromMovie algorithm with different hyperparameter sets. The coefficients for EV, BB, and SC were kept constant at 10^3^, while coefficient for FF was drawn randomly on a logarithmic scale between 0 and 10^6^. For smaller values of *k*_*F F*_ the model should generally behave as an independent modeling strategy, similar to how most models operate by treating each single-cell contact map separately.

The results of this analysis are shown in Fig. 2B. Each point on the plot represents a single hierarchical ChromMovie simulation. A Loess curve was fitted to the data, with a zoom-in view provided in the top-left corner. The analysis revealed a clear minimum RMSD around *k*_*F F*_ ≈ 94.1. Notably, at this FF coefficient value, the reconstruction was significantly enhanced compared to the independent modeling with *k*_*F F*_ ≈0. Although this result was obtained from an in silico example, it provides initial evidence that our approach can improve MD chromatin reconstructions. It also suggests that the FF coefficient should be set approximately one order of magnitude smaller than the other force coefficients.

This result was obtained with all force potentials being harmonic, along with linearization at larger distances. However, alternative formulas for force potentials can also be used. Fig. 2C shows example set of forces that can be used. ChromMovie also automatically computes a variety of metrics based on the simulation results, which are summarized in PDF reports. Fig. 2D showcases some of these metrics. These include the structures’ Radius of Gyration (Rg) and the *α* coefficient from the estimated contact probability scales curve *P* (*s*) ∝ *s*^−*α*^ (as defined in, for example, [13]), both of which are computed across all structure frames and simulation time. Other metrics include local loci variability, local violations of SC contacts, mean violations of all four main force field components, as well as general MD metrics such as energy and temperature during the simulation.

We hypothesize that the local violation detected by ChromMovie (Fig. 2D bottom-right corner) could help identify loci with problematic local topology and knotting. This could also be useful in studies exploring the role of topoisomerases in organizing mitotic chromosomes [81]. However, further investigation is required.

The validity of gradually increasing force coefficients was also tested. In this study, we performed 1, 000 hierarchical ChromMovie simulations, with force coefficients either gradually increasing or remaining constant. The results of this analysis are presented in Fig. 2E. We successfully confirmed that applying a gradual increase of the EV coefficient during the simulation significantly improved reconstruction reliability, as measured by the sum of single-cell contact violations (distances between the 3D simulation results and the upper bound for the flat-bottom of the SC potential). This finding supports similar claims made in previous studies [27, 41], where adding a gradual increase of the EV coefficient (“EV” group) significantly reduced contact violations compared to constant coefficients (“constant” group). However, the effectiveness of this approach was dependent on the specific set of force hyperparameters, and the gradual increase did not consistently enhance the simulation results. Moreover, we were unable to validate the efficacy of a gradual increase for other forces, despite its application in certain studies [41]. We conclude that, in the case of ChromMovie, the overall success of this approach may be context-dependent and is not guaranteed to improve every simulation scenario.

### Benchmarking on ChAIR dataset

ChAIR [80] is a high-throughput method that simultaneously measures gene expression, chromatin accessibility, and chromatin interactions in single cells. By leveraging a microfluidic platform, it can analyze 6, 000–8, 000 cells in a single reaction, providing insights into the 3D epigenomic mechanisms that regulate transcription. Cells with similar transcriptomic profiles during the cell cycle or maturation process were grouped into metacells with same amount of chromatin contacts to increase data volume and reduce stochasticity (see Fig. 3A and B). This dataset was selected to validate our model using real single-cell contact data with specific information about a particular cellular process, namely, cell cycle progression. The dataset comprised 102 metacells, representing the progression through the interphase trajectory phases of the cell cycle, specifically G1, S, and G2M.

**Fig. 3.**
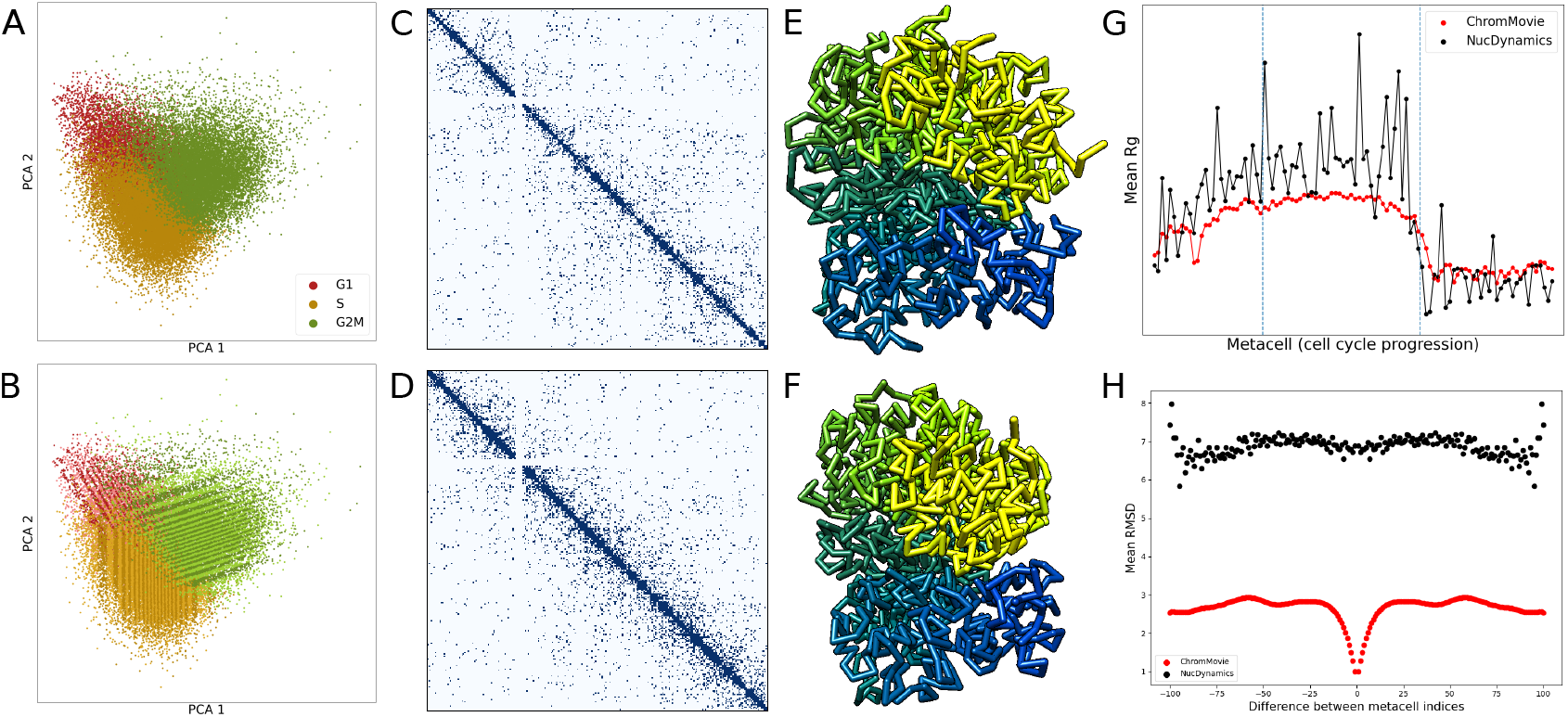
Validation of ChromMovie using ChAIR data. (A) Visualization of single-cell ChAIR data for the k562 cell line using 2D principal component analysis (PCA) (see [80]). (B) The same single-cell data with metacells highlighted as alternating stripes to indicate grouping. (C) and (D) Representative single-cell contact maps for chromosome 12, illustrating a more relaxed chromatin state during the S phase (C) and a more condensed state in the G2M phase (D). (E) and (F) Corresponding 3D chromatin structures generated by ChromMovie for the contact maps shown in (C) and (D), respectively. (G) The mean radius of gyration across all 102 metacells throughout cell cycle progression, comparing ChromMovie and NucDynamics. (H) Mean RMSD values plotted as a function of frame shifts, assessing structural consistency between consecutive frames in both ChromMovie and NucDynamics models.

Based on the contact data derived from these metacells (see Fig. 3C and D), we generated three-dimensional chromatin structure models using both ChromMovie and NucDynamics. For both algorithms, hierarchical modeling was applied with a consistent final resolution of 100kb. Our analysis revealed that both methods were capable of reconstructing the chromosome compaction pattern observed during cell cycle progression, as described in previous studies [65], from early G1 to late G2m phase (Fig. 3G). Chromatin compaction was quantified using the mean radius of gyration of overlapping structural bead segments. Notably, while both methods successfully captured cell cycle dynamics, the structures produced by ChromMovie exhibited considerably lower variability, which aligns with the expected regularizing effect of the frame forces.

Furthermore, we evaluated whether ChromMovie could not only accurately recreate cell cycle progression dynamics, but also achieve smoother transitions and greater similarity between consecutive frames, thereby producing a more biologically relevant evolution of 3D chromatin structure. To this end, the RMSD was computed for all frame pairs in both ChromMovie and NucDynamics structures. The mean RMSD differences were used to assess the degree of metacell displacement between consecutive frames. This analysis provided insights into the correlation of each frame’s structure within the overall “movie” of cell cycle progression. The results demonstrated that the ChromMovie-generated structures exhibited significantly higher similarity across frames compared to those obtained from the independent NucDynamics simulations, thereby validating the robustness of our approach.

### Spatial modeling of genomic features in single cells

Having confirmed that ChromMovie can successfully reproduce key chromatin dynamics associated with cell cycle progression, as reported in previous studies, and does so in a smoother and more continuous manner, we next aimed to evaluate whether the method can also capture other established features of nuclear chromatin compaction. To this end, we tested ChromMovie using diploid cell maturation contact data from [80], derived from vascular mouse brain tissue. The mouse cells were analyzed at 2, 11, 95, 365 and 720 days, resulting in five distinct frames for the ChromMovie simulation. Since the original dataset did not exclusively represent a single cell type, we subsampled the contact data to 5, 000 contacts per frame to ensure consistency. This dataset allowed us to assess whether ChromMovie could recapitulate well-known phenomena such as the emergence of chromosomal territories within the nucleus and the segregation of chromatin compartments.

Figure 4A shows an example frame of the results of this analysis with distinct chromosomal territories marked by contrasting colors. To check whether the chromosomal territories pattern correspond to those reported by the previous studies we estimated the degree of inter-chromosomal intermingling. We followed the definition of intermingling as defined in [38] as the percentage of chromosome beads surrounded by at least four other beads from a different chromosome within a threshold distance of 2 bead diameters. In our case, the bead diameter can be defined as the optimal distance of the backbone force *d*_*BB*_ described previously.

**Fig. 4.**
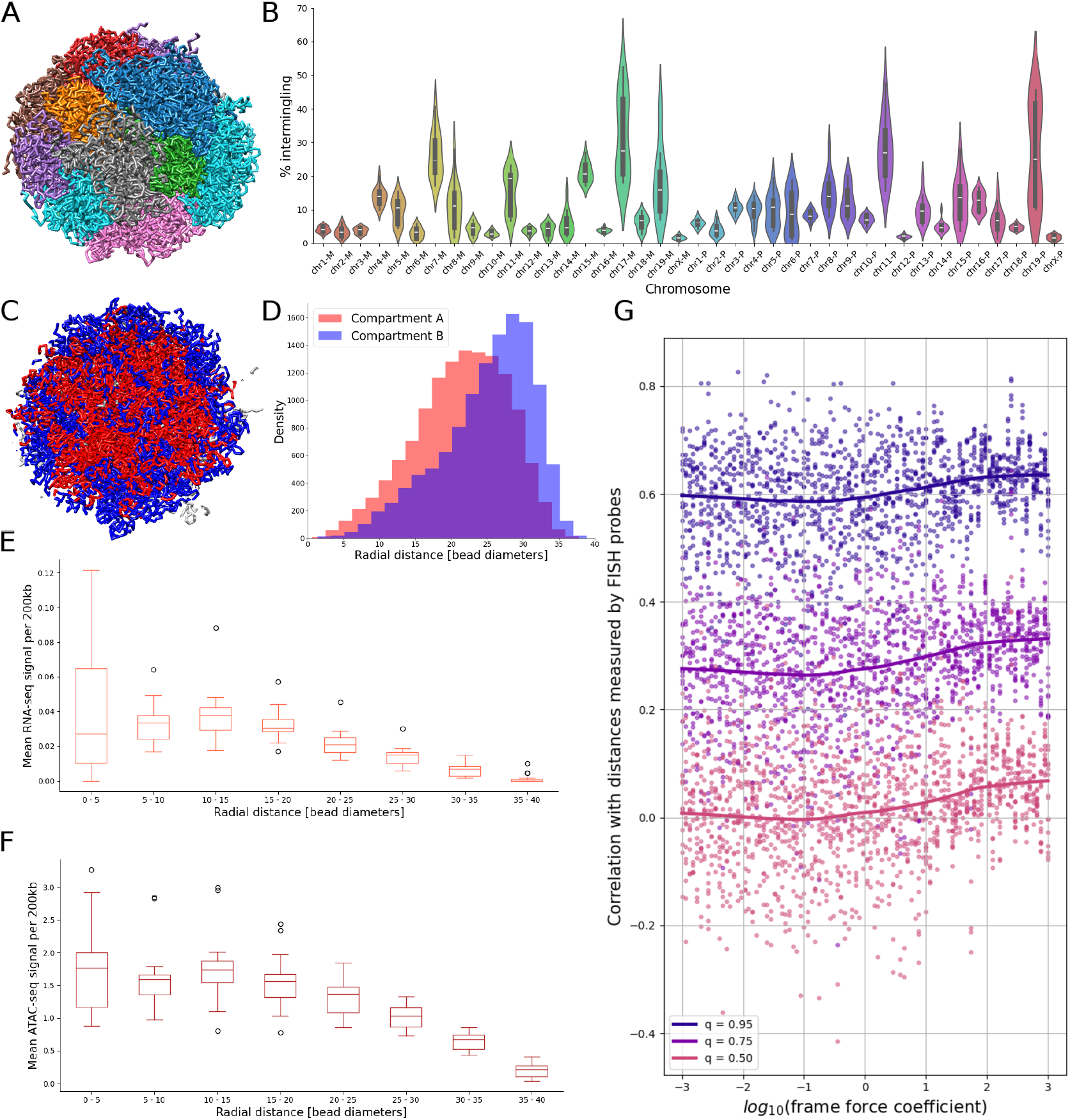
Analysis of ChromMovie-derived structural features using single-cell data. (A) ChromMovie reconstruction of a single-cell, whole-genome diploid mouse nucleus. Individual chromosomes are shown in distinct colors to visualize chromosomal territories. (B) Quantification of inter-chromosomal intermingling based on the structure in (A). Maternal chromosomes are labeled with the suffix “-M” and paternal chromosomes with “-P”. (C) ChromMovie model of the same cell highlighting chromatin compartmentalization. Compartment A (red) and compartment B (blue) are visualized in a central nuclear cross-section. Grey segments represent regions with unknown compartment (mostly centromeres and telomeres). (D) Radial distribution of compartments A and B based on the model in (C), demonstrating preferential localization of active (A) and inactive (B) regions within the nuclear volume. (E) Radial distribution of RNA-seq signal from the ChAIR dataset, showing that transcriptionally active regions are enriched closer to the nuclear center. (F) Radial distribution of ATAC-seq signal from the same dataset, indicating increased chromatin accessibility near the nuclear interior. (G) Validation of ChromMovie structural models using 3D-FISH probe data from Beagrie et al. (2017) [19]. Each point represents a unique combination of cell, frame force coefficient, and the corresponding quantile (0.50, 0.75, or 0.95) of the correlation distribution between modeled and experimental distances. Loess trend lines are shown for each quantile to highlight the effect of frame force regularization.

The results of this analysis are summarized in Figure 4B. We observed intermingling values ranging from approximately 5% to 20%, consistent with previously reported ranges [27,38]. However, we note that some chromosomes in our diploid model exhibited a comparatively higher degree of intermingling. Using the same simulation, we also demonstrated that ChromMovie is capable of capturing key features of compartment segregation. Specifically, we observed that the transcriptionally active compartments A preferentially localize toward the nuclear center, while the B compartments tend to occupy the peripheral regions of the nucleus (Figure 4C and D).

Additionally, ChAIR data provide a unique opportunity to simultaneously analyze single-cell contact data along with transcriptomic (RNA-seq) and chromatin accessibility (ATAC-seq) information. To assess whether our computational framework can generate meaningful spatial distributions for these genomic features, we conducted an integrative analysis using a subset of 15 Patski mouse cells. These cells were selected to meet the following criteria: (1) each contained at least 6, 000 chromatin contacts; (2) all three interphase stages (G1, S, and G2/M) were equally represented with five cells each; and (3) the selected cells formed a closed ring of consecutive points on the PCA plot in Figure 3 A, representing a continuous trajectory through the cell cycle.

We performed a ChromMovie simulation at 200*kb* resolution using 15 frames, each corresponding to one of the selected cells, thereby modeling a full cycle of cell progression. For each simulation frame, we computed the average number of single-cell RNA-seq reads mapped to chromatin beads as a function of their radial distance from the center of mass of the nucleus. The same analysis was repeated for the ATAC-seq signal.

We observed that both RNA-seq and ATAC-seq signals were most concentrated at close to intermediate radial distances from the nuclear center (5 − 20 bead diameters), with the signal intensity gradually decreasing at larger distances (Figure 4E and F). These observations align with established biological knowledge: the nuclear periphery is typically associated with densely packed, transcriptionally inactive heterochromatin, whereas the nuclear interior is enriched with open, transcriptionally active euchromatin. Consequently, both higher ATAC-seq signal and RNA-seq read density are expected to localize closer to the nucleus center, reflecting regions of increased chromatin accessibility and active gene expression.

### Validation with 3D-FISH

Following the example of several previous studies [36, 38, 39], we used the 3D-FISH dataset from Beagrie et al. (2017) [19] as an independent means of validating our methodology. This dataset includes eight pairs of fluorescent probes located on chromosomes 3 and 11, with each pair associated with a set of spatial distance measurements. For validation purposes, the median value of these distances was used, serving as a ground truth reference for comparison with our modeled chromatin structures. Although 3D-FISH data were obtained from mouse embryonic stem cells (mES), whereas contact data were derived from the Patski mouse fibroblast cell line, both datasets used for this validation originate from the mouse embryo.

We performed this validation using the same 15 mouse cells representing cell cycle progression as described in the previous section. Prior studies have commonly assessed model accuracy by computing the Pearson correlation between the median 3D-FISH distances and the corresponding Euclidean distances derived from computationally reconstructed chromatin structures. However, since our data are diploid, we adapted this procedure to account for the presence of both maternal and paternal chromosome copies. Rather than computing a single Pearson correlation, we evaluated all possible pairwise combinations of maternal and paternal chromosome copies for each probe pair. For each combination, we computed a Pearson correlation, resulting in a distribution of correlation values for each probe pair.

From these distributions, we extracted quantiles at the 0.5, 0.75, and 0.95 levels to assess model performance. While the specific chromosome pairing observed in the 3D-FISH experiment remains unknown, a more accurate chromatin model should exhibit increased values across these correlation quantiles, indicating stronger concordance with the experimentally derived spatial constraints.

We employed this approach to validate the core concept underlying ChromMovie, namely, the inclusion of an additional force that connects consecutive frames in the simulation. Using the aforementioned 15 mouse cells, we performed ChromMovie simulations across 100 different values of the frame force coefficient, ranging from 10^−3^ to 10^3^, to determine whether this parameter could improve the concordance with 3D-FISH data and, therefore, enhance the accuracy of chromatin structure reconstruction.

Figure 4G presents the three quantiles (*q* = 0.5, *q* = 0.75, *q* = 0.95) of the correlation distributions between modeled distances and the experimental FISH probe distances, plotted against varying frame force coefficients. Across all quantiles, we observed that low frame force values—where the force is effectively negligible—consistently resulted in weaker correlations with the FISH data. In contrast, higher frame force coefficients yielded stronger correlations, with the trend plateauing around 10^2^, which is a value comparable to the coefficients of other force field components. To assess statistical significance, we performed Wilcoxon rank-sum tests comparing quantile values for simulations with *k*_*F F*_ *<* 1 versus those with *k*_*F F*_ ≥ 1. All quantiles yielded highly significant p-values (*p <* 10^−10^), confirming that the inclusion of the frame force significantly improves model performance.

These results support the conclusion that applying temporal regularization across single-cell simulations, by linking frames, can meaningfully enhance the fidelity of chromatin structure reconstruction, as measured by agreement with independent 3D-FISH data.

### Computational time

We evaluated the computational time for the GPU simulation of the whole genome ChromMovie simulation of the diploid mouse genome from [80]. Figure 5 shows the results of this analysis with respect to the number of cells used for the simulation and a number of resolutions: 5*Mb*, 2*Mb*, 500*kb* and 200*kb*. All cells were selected so that their total number of contacts was around 2000(±30). The calculations were performed on a single NVIDIA A100 GPU. We note that on average 2-fold increase of the simulation resolution and 2-fold increase in the number of cells in the simulation result in ∼ 4-fold and ∼ 2.5-fold increase in computational time, respectively.

**Fig. 5.**
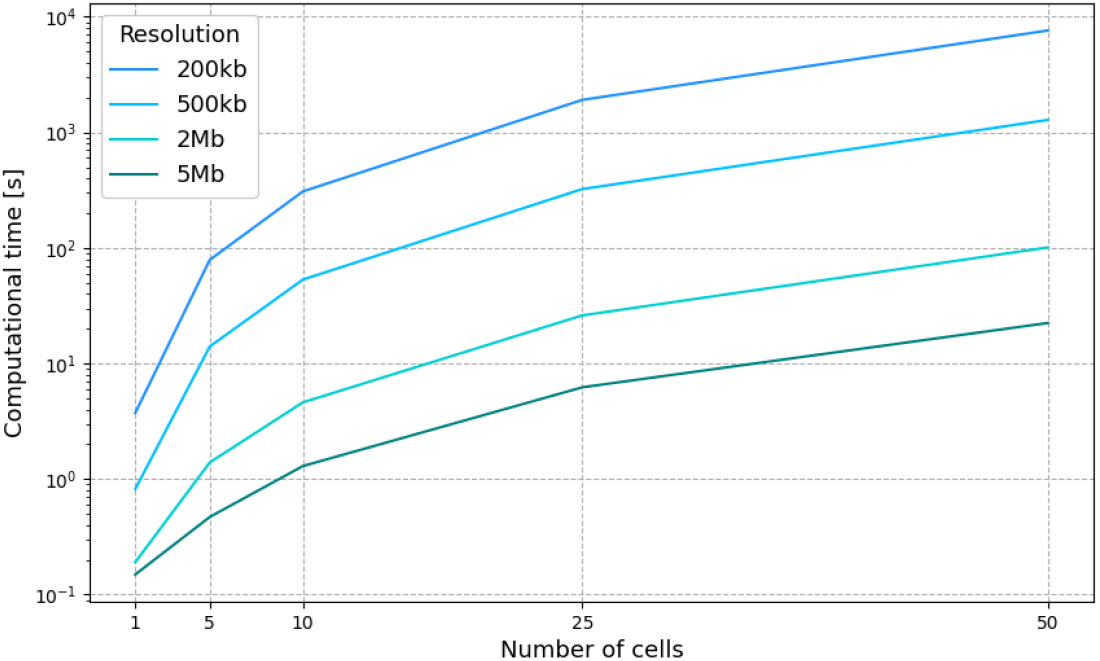
Benchmarking of ChromMovie computational time on a single NVIDIA A100 GPU. For each number of cells a multi-resolution simulation was conducted for resolutions 5*Mb*, 2*Mb*, 500*kb* and 200*kb*.

## Discussion

The challenge of reconstructing chromatin folding at the chromosome level is a critical issue in biology and bioinformatics [82]. Significant attention has been given to deciphering the 3D chromatin structure at single-cell resolution, which reveals patterns related to the cell cycle and cell maturation [65]. In this work, we present ChromMovie: a comprehensive tool designed for studying 3D chromatin structures at the chromosome level throughout any linear cellular process, such as the cell cycle or cell maturation.

While some methods, like HSA [83], integrate multiple Hi-C data tracks, they primarily focus on integrating data from different restriction enzymes, rather than incorporating single-cell information within the context of a specific cellular process. This focus on cellular process context is a core innovation of ChromMovie, and we consider it a novel contribution to the field. We believe that our software will be valuable for researchers studying the changes in 3D chromatin structure during various cellular processes.

We demonstrated that the frame force, a novel feature of our method compared to other approaches, improves the quality of chromatin structure reconstruction, both in the in silico studies and in real single-cell data. Additionally, we showed that ChromMovie effectively recreates chromosome compaction during cell cycle progression in a manner comparable to existing methods. Furthermore, we demonstrated that ChromMovie achieves reduced variance in single-cell structures, potentially capturing more biologically meaningful chromatin conformational changes.

In creating ChromMovie, we drew inspiration from various methods for 3D chromatin structure reconstruction that were developed previously [42]. A particularly influential model for us was NucDynamics [27], which we emulated in several ways. In our algorithm, we aimed to incorporate the best practices from earlier models that provided valuable insights into single-cell chromatin dynamics. These practices include hierarchical modeling, manipulation of the excluded volume force, contact force proportional to the number of contacts in a bin, and the use of different formulas for attractive forces with linearization for long distances. ChromMovie represents an attempt to combine these established practices from the scHi-C modeling field into a novel idea of simultaneous scHi-C modeling.

We also believe that both the fields of 3D chromatin modeling from single-cell Hi-C data (or from 3D contact data in general) and cellular trajectory inference methods can benefit from the development of joint models that leverage the strengths of both fields. To the best of our knowledge, ChromMovie is the first attempt at such a fusion. By developing ChromMovie, we aim to contribute to a more holistic approach to modeling chromatin changes in the context of the specific cellular processes they undergo. Finally, ChromMovie was designed to work with single-cell Hi-C data that have already been ordered in a linear fashion. While this linear assumption is suitable for modeling processes like cell maturation, it may not capture the full scope of the spatio-temporal relationships between cells engaged in various cellular processes. We believe that, in the future, methods that incorporate a broader range of cellular trajectory topologies will be valuable to researchers studying chromatin structure dynamics at the single-cell level.

## Data and code availability

The source code of the method is publicly available on Github: https://github.com/SFGLab/ChromMovie.

The compartments used in this study were taken from the 4D Nucleome Data Portal (https://data.4dnucleome.org/) and are freely accessible under the accession code 4DNFI24OBK5V.

Visualizations of in silico and ChromMovie structures in Figure 1 A, C, D, E, Figure 2 A, Figure 3 E, F and Figure 4 A, C were performed using the UCSF Chimera software [84].

## Author contributions statement

K.H.B. and D.P. conceived the idea of the study. K.H.B. performed the experiments, implemented the method, and conducted the experiments. K.H.B. and D.P. analyzed the results. K.H.B., H.C., Y.R. and D.P. wrote and reviewed the manuscript.

## Acknowledgment

Research was funded by Warsaw University of Technology within the Excellence Initiative: Research University (IDUB) programme. This work has been supported by Polish National Science Centre (2020/37/B/NZ2/03757). Computations were performed thanks to the Laboratory of Bioinformatics and Computational Genomics, Faculty of Mathematics and Information Science, Warsaw University of Technology using Artificial Intelligence HPC platform financed by Polish Ministry of Science and Higher Education (decision no. 7054/IA/SP/2020 of 2020-08-28). The work was co-supported by National Institute of Health USA 4DNucleome grant 1U54DK107967-01 and “Nucleome Positioning System for Spatiotemporal Genome Organization and Regulation”.

